# Gestational physical exercise prevents early-life behavioral impairments and normalizes dopamine D2 receptor expression in the frontal cortex in the offspring of a rat model of attention deficit hyperactivity disorder

**DOI:** 10.1101/2023.07.24.550350

**Authors:** Andréa Tosta, Ariene S. Fonseca, Débora Messender, Priscila Siqueira, Sérgio T. Ferreira, Mychael V. Lourenco, Pablo Pandolfo

## Abstract

Attention deficit hyperactivity disorder (ADHD) is characterized by inattention, hyperactivity and impulsivity, and develops most frequently during childhood and adolescence. Spontaneously hypertensive rats (SHR) are the most used experimental model for the study of ADHD. SHR exhibit behavioral impairments that recapitulate phenotypes observed in individuals with ADHD. SHR further develop dopaminergic hypofunction in frontostriatal circuits and an imbalance in dopamine and norepinephrine systems. Maternal physical exercise (e.g., swimming) during pregnancy has been shown to promote angiogenesis, neurogenesis, learning, and memory in the offspring of control rats. We investigated the impact of gestational swimming on behavioral and dopaminergic parameters in childhood (1-2 weeks of age) and adolescent (4-5 weeks of age) SHR and Wistar Kyoto rats (WKY), used as a control. Maternal gestational swimming resulted in a reversal of neurodevelopmental impairments in behavior, assessed by the righting reflex and olfactory recognition tests, in the offspring. Furthermore, during adolescence, SHRs from exercised dams exhibited reduced novelty seeking, an important behavioral trait in this developmental period. Finally, SHRs exhibited increased expression of dopamine transporter (DAT) and D2 receptors (D2R) in the frontal cortex. D2R expression was normalized in the frontal cortex of adolescent SHRs whose mothers were exercised. Results suggest that physical exercise during pregnancy could be an effective preventative strategy against ADHD-associated behavioral and neurochemical phenotypes in the offspring.

**Graphical Abstract:** 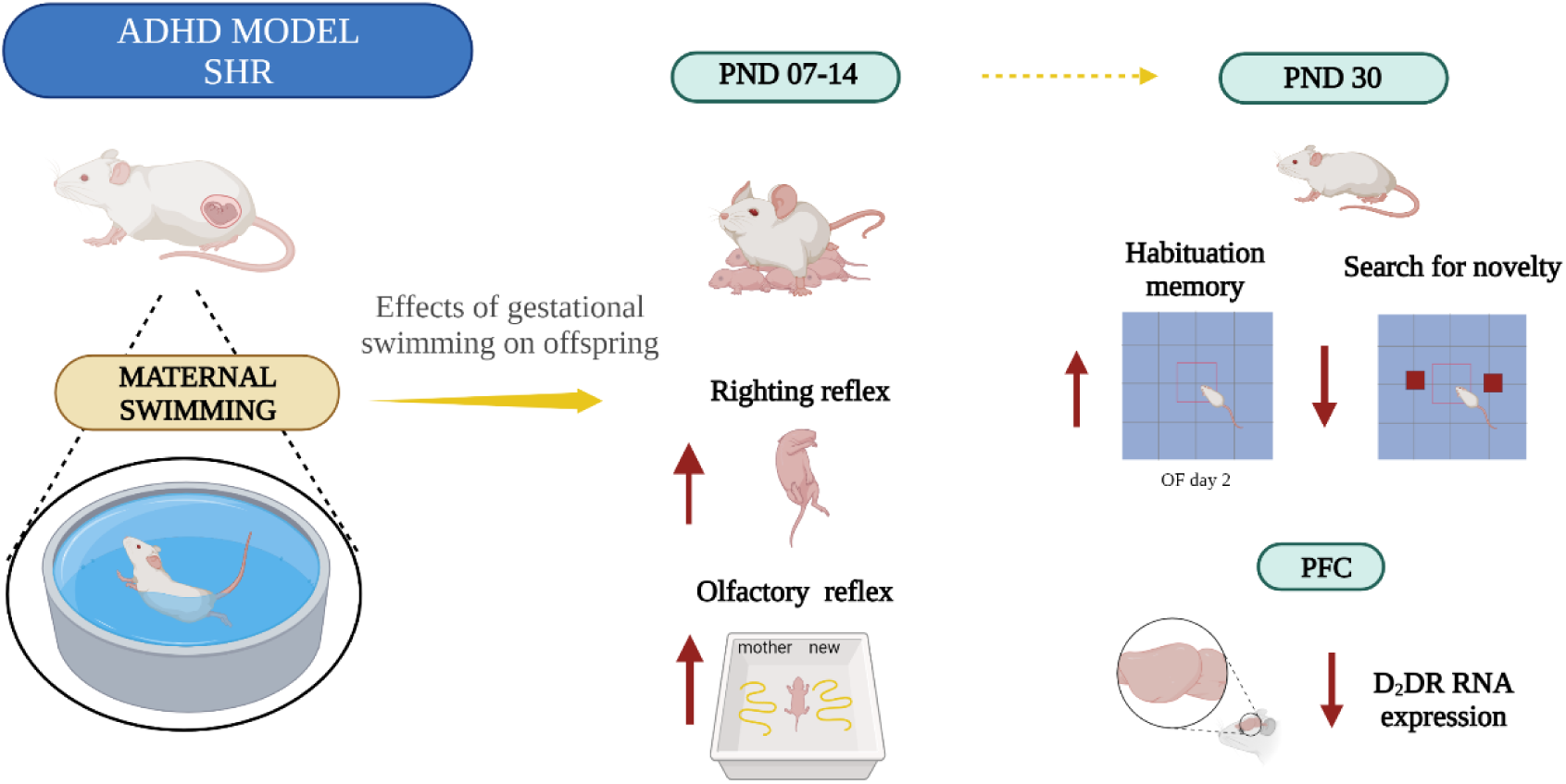

## 1. Introduction

ADHD is a psychiatric disorder characterized by inattention, hyperactivity, and impulsivity (American Psychiatric Association, 2013) that usually develops in childhood and adolescence (Faraone, 2018; Faraone et al., 2015; Polanczyk and Rohde, 2007). While considerable advances have been made in the pathophysiology of ADHD, a complete understanding of its risk factors and options for symptom management is still lacking.

Spontaneously hypertensive rats (SHR) are the most used experimental model for the study of ADHD, when compared to Wistar Kyoto rats (WKY). (Sagvolden, 2000a). SHR exhibit behavioral traits observed in individuals with ADHD (Russell et al., 2006; Sagvolden et al., 2009, 1992; Sagvolden and Sergeant, 1998), in addition to dopaminergic hypofunction in frontostriatal circuits and imbalances between the dopaminergic and norepinephrinergic systems. SHR respond positively to methylphenidate (MPH), a drug that inhibits dopamine transporters and is used as treatment for human ADHD (Russell, 2011).

Lifestyle factors during pregnancy, such as physical exercise or drug abuse, affect development and neural plasticity (Nelson et al., 2011). Exercise has been reported to promote numerous benefits for overall health and brain function. Studies in rodents and humans revealed that regular physical exercise has positive effects on brain function by improving memory, learning, angiogenesis, neurogenesis, and neurotransmitter release, in addition to inducing neuroprotection against injuries (Favero et al., 1993; Gómez-Pinilla et al., 2002; Isaac et al., 2021; Lourenco et al., 2019; Molteni et al., 2004, 2002; Noakes et al., 2005; Sanches et al., 2021, 2018, 2017; Swain et al., 2003; Praag, 2008; Praag et al., 2005) Exercise also increases the synthesis, release, and availability of dopamine in the brain (Cho et al., 2014; Yoon et al., 2007). In adolescent SHR, swimming improves cognition, increases protein levels of tyrosine hydroxylase (TH), the rate-limiting enzyme in dopamine biosynthesis, and rescues reduced dopamine D2 receptor (D2R) expression (Ko et al., 2013).

Mild-intensity exercise during pregnancy has been considered safe for both mother and child (Kim et al., 2007). Gestational exercise promotes muscle strength and endurance,and alleviates pregnancy-related anxiety and depression (Amorim et al., 2007; Bungum et al., 2000; Kim et al., 2007; Polley et al., 2002). Exercise during pregnancy has also been reported to provide benefits to offspring, such as increased hippocampal BDNF mRNA, neurogenesis, and improved spatial learning (Parnpiansil et al., 2003) (Marcelino et al., 2016, 2015, 2013; Sanches et al., 2021b, 2017b). Here we investigated the effects of swimming during pregnancy in the offspring of an ADHD animal model, particularly on sensorimotor reflexes, locomotor activity, emotional behaviors, and on dopaminergic parameters in the prefrontal cortex.

## 2 Material and methods

### 2.1 Experimental subjects

Male and female isogenic rats of approximately 90 days old (WKY and SHRs) were used for mating. They were obtained from the Multidisciplinary Center for Biological Research (CEMIB) of the State University of Campinas (UNICAMP) (Campinas, SP, Brazil) and kept in the sectoral vivarium of the Department of Neurobiology, Institute of Biology of the Fluminense Federal University (Niterói, RJ, Brazil). They were maintained with standard food and water ad libitum, and a 12-h light and dark cycle in air-conditioned rooms (24°C ± 2°C). WKY and SHR rats were housed for mating in Plexiglass boxes (42×34×18 cm), divided as follows: two females and one male per box during mating and 6 per box after weaning of offspring. All animal experiments were performed in accordance with the principles of the “3Rs” and the National Institutes of Health Guide for the care and use of laboratory animals.

### 2.2 Experimental design

After confirmation of pregnancy, rats were immediately subjected to the swimming protocol, from gestational day 0 (GD0) to GD20. Swimming was performed every day of the week, except Saturday and Sunday, and each session lasted 20 minutes. On the twentieth day of gestation, females were placed in isolated cages and remained there until parturition. On post-natal day (PND) 7 and PND 14, offspring were weighed, and neonatal neurodevelopmental tests were performed to analyze development and sensorimotor reflexes. On PND 21, the animals were weighed, weaned, and placed in cages according to sex, strain and treatment. Behavioral tasks were performed during adolescence: elevated plus maze (EPM) (PND 30), open field (OF) day 1 (PND 31), OF day 2 (PND 32), and novelty seeking (PND 33). Twenty-four hours after the last behavioral task (PND 34), the frontal cortex collected for biochemical analysis (Fig. 1).

**Figure 1.**
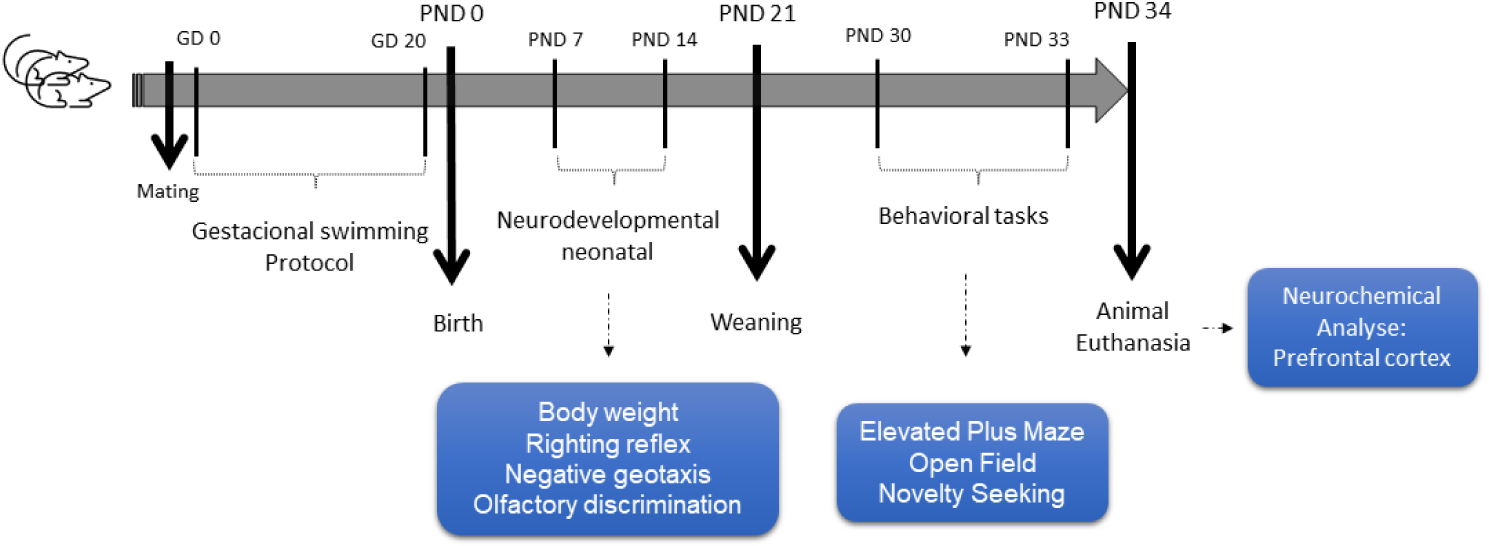
Schematic overview of the experimental design. Male and female rats (90 day old) of strains WKY and SHR mated. After confirmation of pregnancy, rats were immediately submitted to the swimming protocol. The swimming protocol was performed from gestational day 0 (GD0) to G20. Each daily session had 20 minutes. On the twentieth day of gestation, females were housed in isolated cages and remained there until parturition. The day of birth was considered the postnatal day 0 (PND0). At PND 7 and PND 14, animals were weighed and SHIRPA tests were performed to analyze the development and sensorimotor reflexes of the offspring. After PND 21, animals were weaned and housed in cages according to sex and strain. From PND 30, behavioral tasks were performed during adolescence: Elevated Plus Maze (EPM) (PND 30), open field (OF) day 1 (PND 31), OF day 2 (PND 32) and novelty seeking (PND 33). Twenty-four hours after the last behavioral task (PND 34), prefrontal cortex collected for biochemical analysis.

### 2.3 Exercise protocol (swimming)

The protocol for gestational swimming was adopted from Lee et al. (2006). Swimming was performed daily starting at GD0, the day of pregnancy confirmation, until GD20. Female animals were split into 4 groups: 2 control groups (sedentary groups, WKY SED and SHR SED) and 2 exercised groups (swimming groups, WKY SWIM and SHR SWIM). Animals in the WKY SWIM and SHR SWIM groups were individually exposed to a 50 x 30 cm plastic tank with 35 cm-high water heated to a temperature of approximately 30-32°C for 20 minutes / 5 times *per* week. Mothers in the control group, WKY SED and SHR SED, were exposed only to the training environment: a tank filled with water at a depth of approximately 1 cm for 5 minutes. Pregnant rats were weighed every week for pregnancy control. All rats (SED and SWIM) were gently placed in their respective boxes for each session, dried with a towel and returned to their home cage.

### 2.4. Neurological Reflex Testing

On PND 7 and PND 14, sensorimotor reflex tests were performed to examine the effects of gestational swimming on the neurodevelopment of the offspring.

#### Righting reflex

The total time required for the animal to straighten up in a maximum of 60 s was assessed. To do this, the animal was placed in dorsal decubitus and restrained for 5 seconds, and the latency it takes to turn to its original position (ventral decubitus) was measured (Ferguson et al., 2003a; Sanches et al., 2017).

#### Negative geotaxis

The duration of the animal’s coordination reflex was assessed. The ability to return to the opposite head direction in an apparatus with a 30° tilt was assessed by turning 180°. The latency is measured in a maximum time of 60 s (Ferguson et al., 2003b; Sanches et al., 2017).

#### Olfactory recognition

The olfactory recognition test (OR) is a sensory reflex test that measures the time it takes the animal to orient to the odor of the mother. The test was performed in a transparent, rectangular acrylic box containing clean sawdust on one side and sawdust from the original cage impregnated with the odor of the mother on the other side. The animal’s ability to orient to the original sawdust was assessed in a maximum time of 180 s (Ferguson et al., 2003a; Sanches et al., 2017).

### 2.5. Behavioral tasking in adolescence

Animals were acclimatized to the experimental room for 1 hour for both the neurodevelopmental and behavioral tests, which were conducted in a closed, silent room with low-intensity indirect lighting. Equipment and objects were cleaned with 10% alcohol between experiments to avoid odor traces. Animal behavior was recorded with a camera mounted above the apparatus and quantified using AnyMaze® software.

#### Elevated plus maze (EPM)

EPM test was performed in a wooden apparatus coated with gray formica, in which two open arms (50 x 10 cm) and two closed arms (50 x 10 x 45 cm) are connected by a central platform (10 x 10 cm) elevated to a height of approximately 50 cm. Rats were placed on the central platform with their heads facing one of the open arms and observed for 5 min (de Santana Souza et al., 2020; Pellow et al., 1985). The following parameters were recorded: percentage of time (%TA) and entries (%EA) in open arms, and percentage of entries in enclosed arm.

#### Open field (OF)

The OF test was performed in a wooden arena (60 cm^3^) whose walls were covered with white Formica. Rats were placed in the central area and their locomotor activity was recorded for 10 min on day 1 and day 2. On the first day of OF, locomotor activity was assessed in response to a novel environment. The second day of the task assessed habituation memory, a context that is considered familiar (Hall, 1934; Penna et al., 2023; Prut and Belzung, 2003; Walsh and Cummins, 1976).

#### Novelty seeking

On the fourth day of behavioral testing, animals were again exposed to the OF arena with the novelty of having two identical objects in the central area. The time that the animals spent interacting with the objects was recorded for 5 min.

### 2.6. Western blotting

Twenty-four hours after the last behavioral test, the animals were euthanized, and the frontal cortex was removed for neurochemical analyses and mRNA extraction. Cortical samples were stored at -80 °C until homogenization in RIPA buffer (25 mM Tris-HCl pH 7.6, 150 mM NaCl, 1% NP-40, 1% sodium deoxycholate, 0.1% SDS; Thermo Fisher) containing cocktails of protease and phosphatase inhibitors. Samples were resolved in pre-cast 4-20% polyacrylamide gels (BioRad®) with Tris/glycine/SDS buffer and resolved at 100 V for 90 min at room temperature. The gel (30 µg total protein/lane) was electroblotted onto Hybond ECL nitrocellulose using 25 mM Tris, 192 mM glycine, 20% methanol (v/v), pH 8.3, at 350 mA for 60 min at 4°C. Membranes were blocked with 3% BSA in TBS-T for 1 hour at room temperature. The membranes were then incubated under agitation with primary antibodies: anti-DAT, anti-TH_total_, anti-D2DR (1:1,000, diluted in TBS-T containing 3% BSA), anti-β-actin monoclonal antibody (1:2,000, diluted in TBS-T containing 3% BSA) (Supplementary Table 1) overnight at 4 °C. After incubation with secondary anti-rat or anti-mouse IgGs (1:10,000, diluted in TBS-T containing 3% BSA) (Supplementary Table 1) for 120 min, the membranes were washed with TBS-T and developed for imaging in an Odyssey apparatus. The images obtained were analyzed using NIJ Image J to obtain average optical density values of the protein bands.

### 2.7. Quantitative RT-PCR

Total RNA was extracted from frontal cortex samples of adolescent WKY and SHRs using Trizol® (Invitrogen), following manufacturer’s instructions. Isolated total RNA was eluted with 20 µL of nuclease-free water. RNA concentration and purity were determined with BioDrop (Biochrom) by calculating the optical density ratio at 260/280 nm and 260/230 nm. For qRT-PCR, one µg of total RNA was used for cDNA synthesis using the High-Capacity cDNA Reverse Transcription Kit (Applied Biosystems, CA). Quantitative analysis of target gene expression was performed on a 7500 Applied Biosystems real-time PCR system (Foster City, CA) with the Power SYBR Green master mix. β-actin (actb) was used as an endogenous reference gene for data normalization. qRT-PCR was performed in reaction volumes of 12 µL according to manufacturer’s protocols. The primer sequences used for qPCR are described in Supplementary Table 2. Cycle threshold (Ct) values were used to calculate fold changes in gene expression using the 2^-ΔΔCt^ method (Livak and Schmittgen, 2001).

### 2.8. Statistical analyses

All values were expressed as mean ± standard error of the mean (SEM). Two-way analysis of variance (ANOVA) (treatment x strain) was used to analyze the weight of the maternal offspring during pregnancy. Three-way analysis of variance ANOVA (treatment x strain x sex) was performed to analyze offspring weight gain at PND 7 and PND 14, neonatal neurodevelopmental reflexes and all behavioral tests, and biochemical analyzes. For the habituation test, comparing day 1 with day 2 of OF, a three-way ANOVA with repeated measures was used. When results were significant for ANOVA, multiple comparisons were performed using the Tukey *post hoc* test (Tukey test for multiple comparisons). The significance level adopted in all experiments was p ≤ 0.05. Statistical analyzes were performed using JASP software (Jeffreys’s Amazing Statistics Program, Amsterdam) and graphs were generated using Prism 8.0 software (Graphpad®).

## 3 Results

### 3.1 Gestational swimming reduced maternal weight gain

A known benefit of physical exercise to pregnant females is a reduction in net weight gain compared to sedentary mothers (Clapp and Little, 1995). To verify whether swimming during pregnancy affected weight gain in the dams (Fig. 2), a two-way repeated measures ANOVA was performed to compare GD 0 and GD 20. Weight gain was observed for all mothers [F (1,12) = 254.8] (p<0.001). An interaction between strain and treatment [F (1,12) = 9.9] was also observed. Multiple comparisons showed that mothers from the SHR SWIM group had lower weight at the end of pregnancy than the mothers from SHR SED and WKY SED (p<0.01). Thus, gestational swimming selectively attenuated weight gain during pregnancy in SHR.

**Figure 2.**
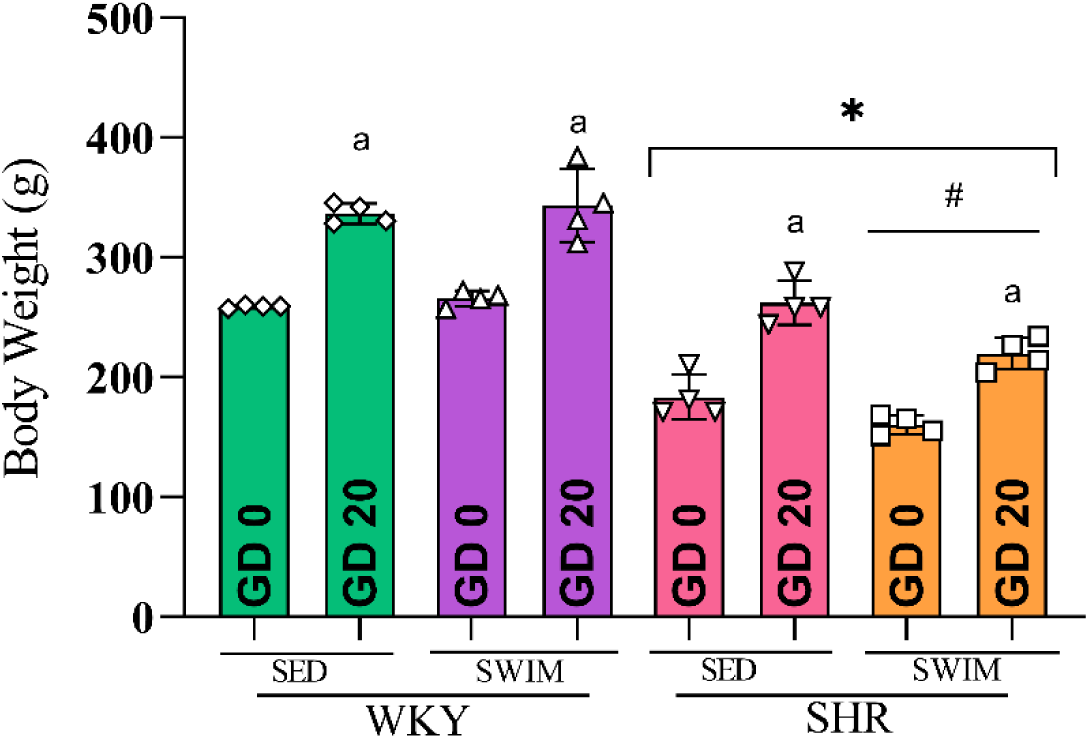
Maternal weight gain during pregnancy. Histogram represent the means ± SEM (n=4-5 animals per group). ^a^p<0.05 difference between day 0 and day 20 (repeated measurements ANOVA; *post hoc* Tukey). *p<0.05 WKY vs. SHR strains, regardless of treatment; ^#^p<0.05 SED vs. SWIM of SHR strain (three-way ANOVA; *post hoc* Tukey).

### 3.2 Swimming during pregnancy decreased the weight of WKY and SHR offspring in PND 07

Weighing of the offspring at PND 7 and PND 14 showed an interaction between weight gain and days of weighing (Fig. 3). A three-way ANOVA analysis was performed for DNP 7 and showed a significant effect for the strain factor [F (1,84) = 89.1] and the strain x treatment interaction [F (1,84) = 19.2]. Multiple comparisons showed that the offspring of WKY animals that performed swimming during pregnancy had lower body mass gain (p < 0.001). On the other hand, the offspring of the SHR SWIM group had a higher body mass compared to the sedentary group WKY (p = 0.01), regardless of sex. At PND 14, differences between strains persisted [F (1,77) = 100.7] and WKY animals continued to have greater body mass than SHRs (p < 0.001).

**Figure 3.**
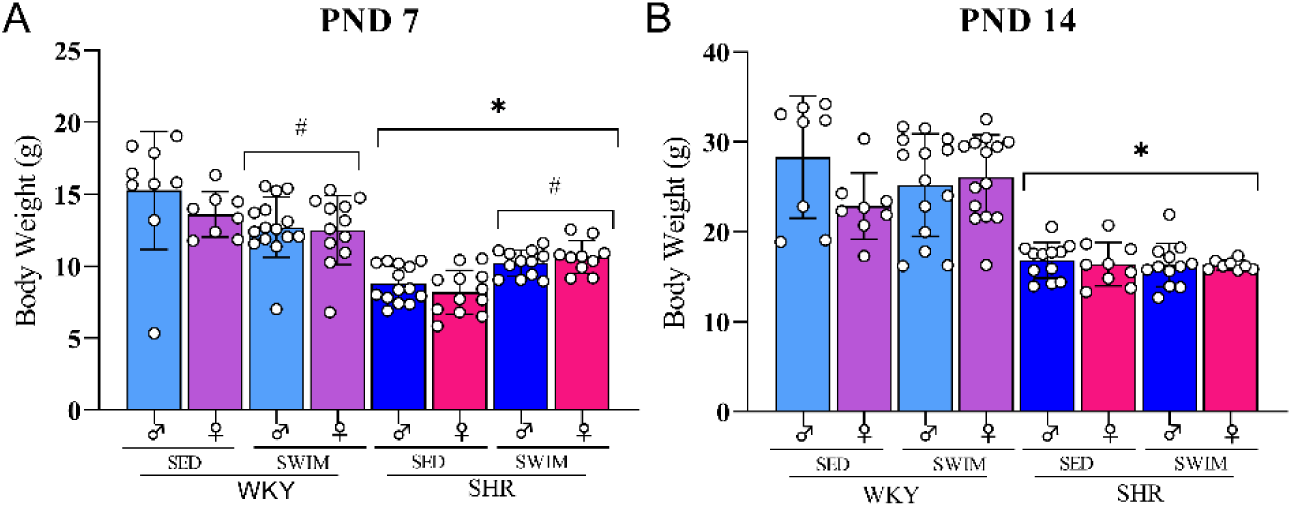
Effects of maternal swimming on offspring weight gain. Comparison between offspring weight gain in PND7 (A, n=9-15 animals per group) and PND14 (B, n=8-14 animals per group). Data are presented as means ± SEM. ^#^p<0.05 SED vs. SWIM treatments in the same strain, regardless of sex; *p<0.05 WKY vs. SHR strains, independent of treatment and sex (three-way ANOVA; *post hoc* Tukey).

### 3.3 Gestational swimming prevented neonatal neurodevelopmental delays in SHR offspring

For the righting reflex, a three-way ANOVA showed a significant difference in the strain x treatment x sex interaction at PND 7 [F (1,80) = 4.5] (Fig. 4). Multiple comparisons revelated that SHRs showed developmental impairment, i.e., they took longer to righting than WKY animals, regardless of sex (p < 0.01). Gestational swimming decreased righting latency in SHR (p < 0.01) (Fig. 4A). On PND 14, analyses revealed a significant interaction between strain x treatment [F (1,78) = 6.3]. SHR offspring also had impairments in the righting reflex compared with the WKY animals (p < 0.001), which was prevented in SHR from exercised mothers (p < 0.001) (Fig. 4B).

**Figure 4.**
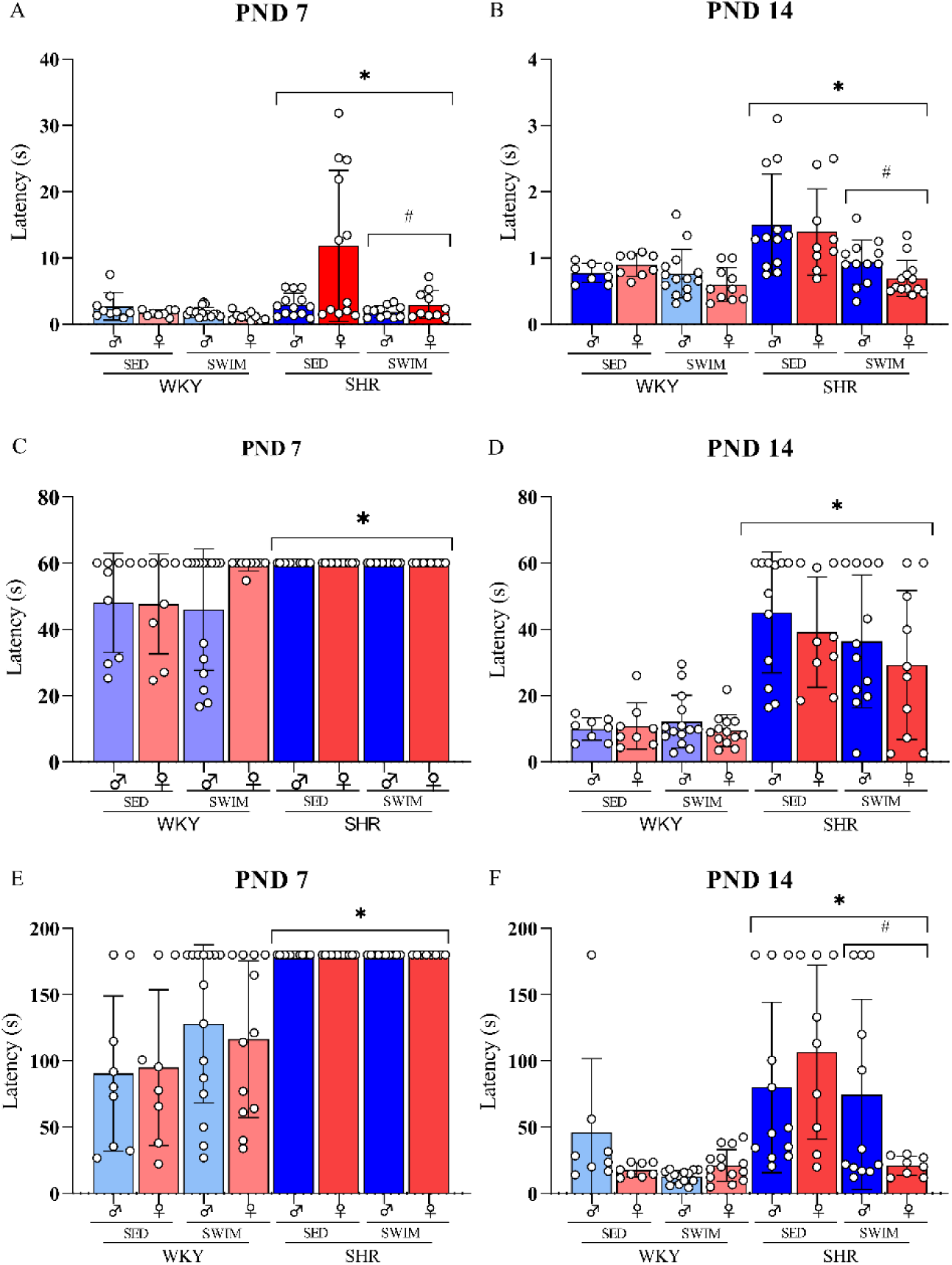
Effects of maternal swimming on neonatal neurodevelopmental reflexes in the offspring. Bars show the means ± SEM of latency in the righting reflex in PND7 (A, n=9-15 animals per group) and PND14 (B, n=8-14 animals per group); in the motor coordination reflex, by the negative geotaxis test, in PND7 (C, n=9-15 animals per group) and PND14 (D, n=8-14 animals per group); in odor discrimination, by the olfactory recognition test, in PND7 (E, n=9-15 animals per group) and PND14 (F, n=8-14 animals per group). *p<0.05 WKY vs. SHR strains, regardless of treatment and sex, ^#^p<0.05 SED vs. SWIM treatments in the same strain, regardless of sex (three-way ANOVA; *post hoc* Tukey).

In the negative geotaxis test, PND 7 showed an effect of strain [F (1,81) = 20.1]. SHR were unable to complete the task and thus reached the maximum required time of 60 seconds, showing impaired motor coordination compared to WKY (p < 0.001) (Fig. 4C). On PND 14, this difference between strains persisted [F (1,78) = 71.1], and SHR showed impaired motor coordination compared to WKY (p < 0.001) (Fig. 4D). Gestational swimming had no effect on the motor coordination reflex of the offspring, regardless of strain and PND (Figs. 4C and 4D).

In the olfactory recognition test on PND 7, three-way ANOVA revealed a significant effect of strain [F (1,80) = 62.8]. SHRs took significantly longer to detect the maternal odor than WKY animals (p < 0.001) (Fig. 4E). On PND 14, a three-way ANOVA reveled a significant strain x treatment x sex interaction [F (1, 77) = 8.3]. SHR displayed decreased olfactory recognition compared to WKY (p < 0.001). Swimming during gestation caused a significant positive effect on sensory development in the offspring, with the SHR SWIM group displaying better olfactory discrimination performance than the SHR SED group (p < 0.01).

### 3.4 Swimming during pregnancy did not alter anxiety-related behavior of adolescent WKY or SHR offspring in the elevated plus maze (EPM)

On PND 30, the EPM was performed to measure anxiety-like behavior. Three-way ANOVA revealed no significant effects in the percentage of entries and time spent in open arms, or in the percentage of entries into enclosed arms (Fig. 5).

**Figure 5.**
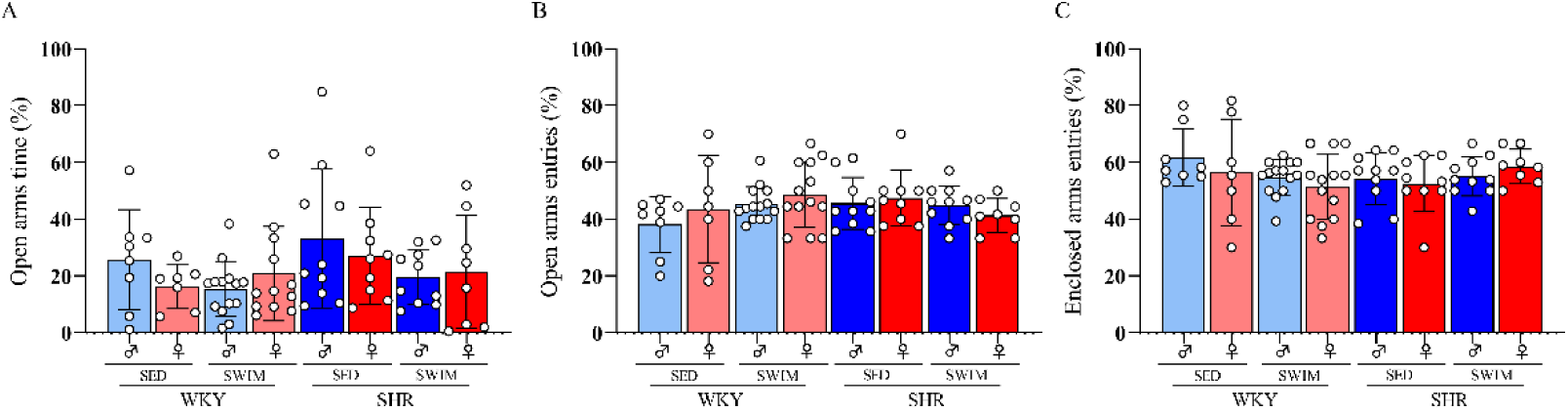
Effects of maternal swimming on anxiety-like behavior in adolescents. Bars represent means ± SEM (n=8-14 animals per group) of percentage of (A) time and (B) entries in the open arms, and of percentage of (C) entries in the enclosed arms.

### 3.5. Effects of swimming during pregnancy on locomotor activity, memory habituation and emotional behavior of WKY and SHR adolescent offspring in open field

For total distance, three-way ANOVA showed a significant effect for the strain x treatment interaction [F (1.75) = 9.3] (Fig. 6?). SHR covered a shorter distance compared to the WKY group (p < 0.05), regardless of treatment or sex. Gestational swimming decreased the total distance traveled by WKY, regardless of sex (p < 0.01) (Fig. 6A). In the first OF session, the following emotional parameters were also recorded in the central zone: latency of first entry (Fig. 6B), percentage of time (Fig. 6C) and locomotion (Fig. 6D). Three-way ANOVA showed a significant effect of strain in latency to enter the central area [F (1,73) = 10.2] and in central locomotor activity [F (1,73) = 12.6]. As expected, multiple comparisons indicated that SHR displayed lower latency to enter (p < 0.01) and higher activity (p < 0.001) in the central area when compared to WKY, regardless sex or treatment.

**Figure 6.**
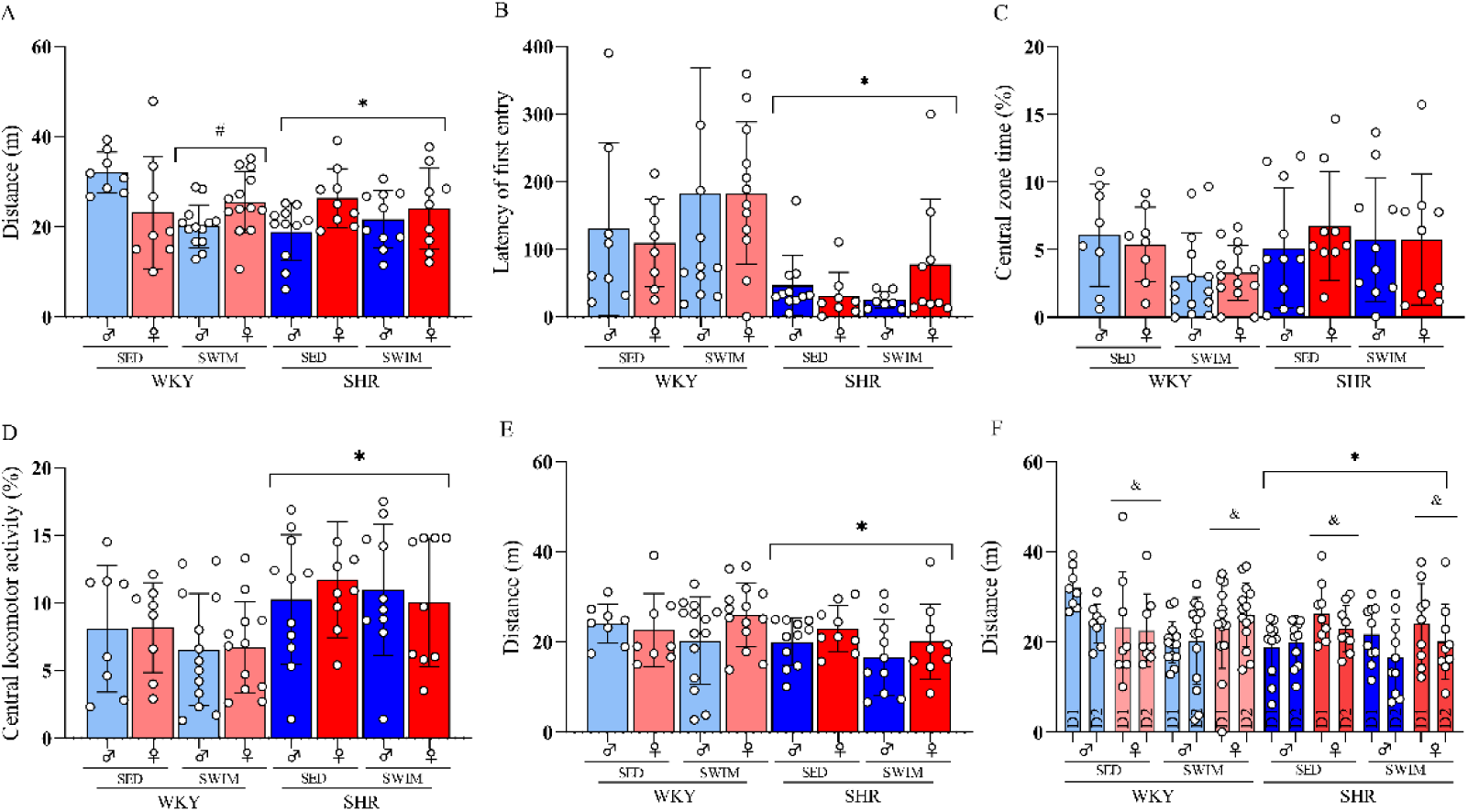
Effects of maternal swimming on locomotor activity, memory habituation and emotional behavior in adolescents. Bars show the mean or median (± SEM or interquartile range) of (A) total distance traveled (m) in the open field arena during the first 10-min test session (day 1). Also in the first session, the following parameters were recorded in the central zone: (B) latency of first entry, (C) percentage of time and (D) locomotion. In the second session (day 2), (E) total distance traveled (m) in the open field during 10 minutes. (F) Total distance traveled (m) in the open field arena during the 1^st^ and 2^nd^ test session. *p<0.05 WKY vs. SHR strains, regardless of treatment and sex, ^#^p<0.05 SED vs. SWIM treatments in the same strain, regardless of sex (three-way ANOVA; *post hoc* Tukey); *p<0.05 WKY vs. SHR strains, regardless of treatment and sex, ^&^p<0.05 male vs. female, regardless of treatment or strain (repeated measures ANOVA; *post hoc* Tukey).

On the second day of OF, a three-way ANOVA revealed a significant effect of the strain [F (1,75) = 4.0] (Fig. 6F). WKY presented a greater distance traveled when compared to SHR (p < 0.05), regardless of treatment or sex. Repeated measures ANOVA showed a significant effect of sex [F (1,73) = 17.8], regardless of treatment, strain, or repetition. The female SHRs presented a decrease in the distance traveled in the 2nd OF session (p < 0.05).

### 3.6. Gestational swimming reduces novelty seeking in adolescent SHR offspring

Three-way ANOVA revelated significant effects of strain [F (1,74) = 15.5] and treatment [F (1,74) = 7.6] (Fig. 7?). Multiple comparisons showed that SHRs explored objects longer than WKYs (p < 0.001). Both adolescent WKY and SHR offspring showed decreased amount of time spent exploring objects (p < 0.01).

**Figure 7.**
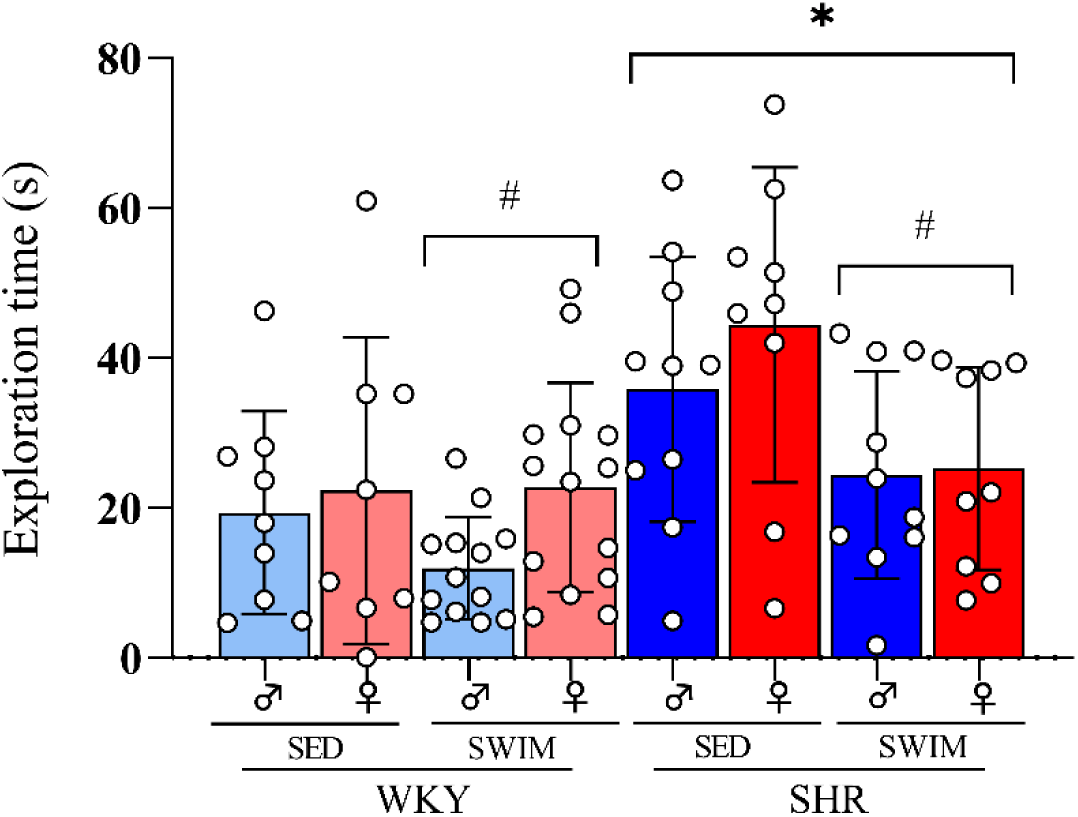
Effects of maternal swimming on novelty seeking in adolescents. Bars show means ± SEM of exploration time of two identical objects placed in the central area of the open field arena (n=8-14 animals per group). *p<0.05 WKY vs. SHR strains, regardless of treatment and sex, ^#^p<0.05 SED vs. SWIM treatments in the same strain, regardless of sex (three-way ANOVA; *post hoc* Tukey).

### 3.7. Swimming during pregnancy does not affect levels of proteins involved in dopaminergic neurotransmission in the frontal cortex (FC)

To investigate whether gestational swimming influenced the dopaminergic system, immunodetection by Western blotting was performed to measure protein levels of TH (Figs. 8A e 8B), DAT (Figs. 8C e 8D), and D2_R_ (Figs. 8E e 8F) in PFC. Three-way ANOVA showed no significant differences in the protein levels examined.

**Figure 8.**
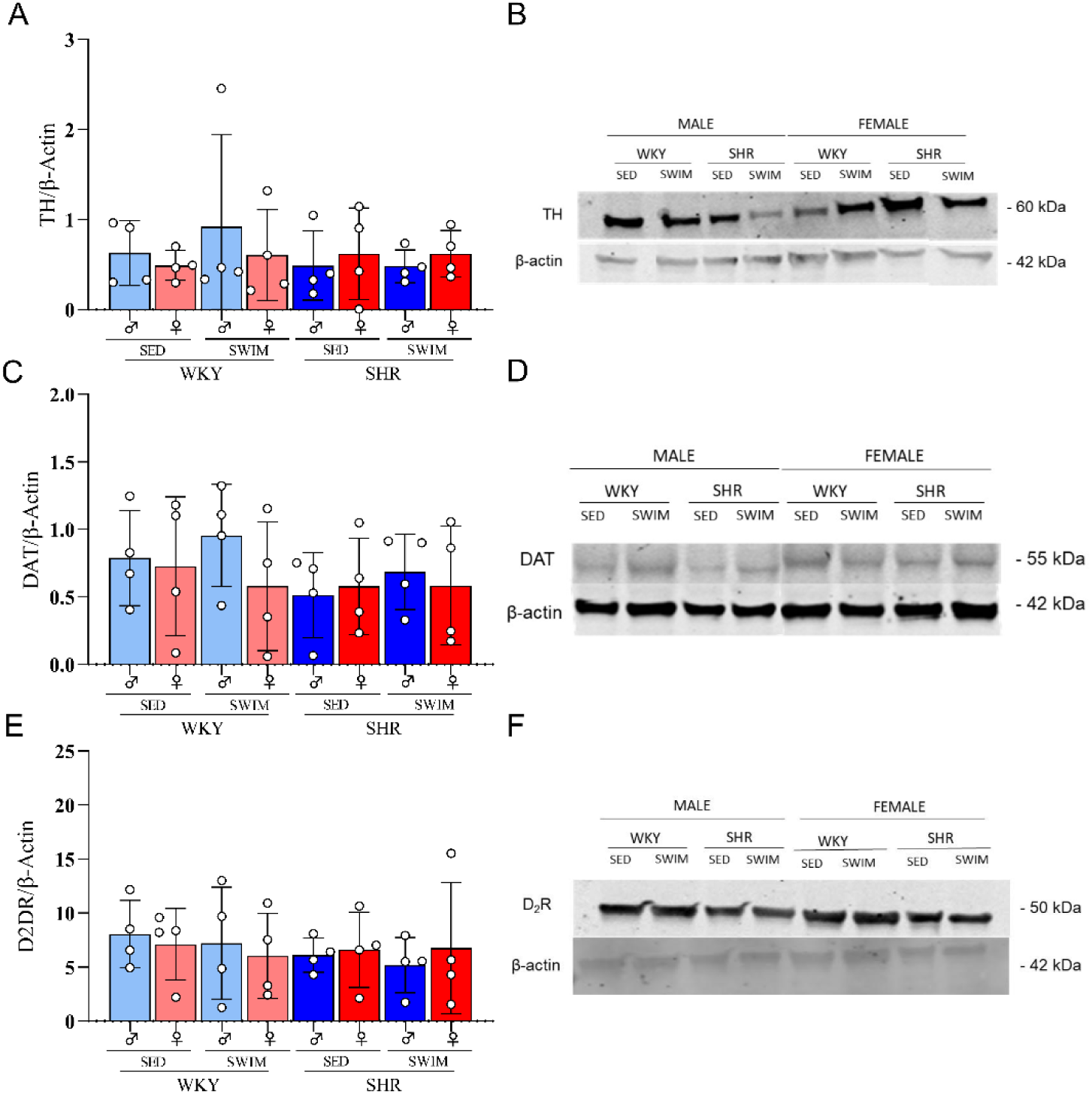
Effects of maternal swimming on density of dopaminergic parameters in prefrontal cortex. Histogram represent the means ± SEM (n=4 animals per group) of density of (A) tyrosine hydroxylase enzyme (TH), (C) dopamine transporters (DAT) and (E) dopaminergic D_2_ receptors (D_2_DR). Representative blots (B, D, F) in samples of the prefrontal cortex of WKY and SHR rats exposed or not to maternal swimming (SWIM and SED).

### 3.8. Gestational swimming decreased the expression of dopamine D2 receptors in the SHR in the PFC

Total RNA expression was analyzed in PFC for TH (Fig. 9A), DAT (Fig. 9B), and D2R (Fig. 9C). No changes in TH expression were detected comparing male or female WKY or SHR rats, whether sedentary or exercised (Fig. 9A). Regarding DAT mRNA content, three-way ANOVA revelated a significant effect of strain [F (1,29) = 7,781]. SHR rats had higher levels of DAT (p < 0.01) compared to control WKY rats, regardless of sex. Regarding D_2_D_R_ expression, analysis showed a significant strain x treatment interaction [F (1,31) = 6.76]. SHRs showed higher D2R expression compared to WKY animals (p < 0.05). Gestational swimming increased D2R expression in WKY (p < 0.05) and decreased in SHR (p < 0.05), regardless of sex.

**Figure 9.**
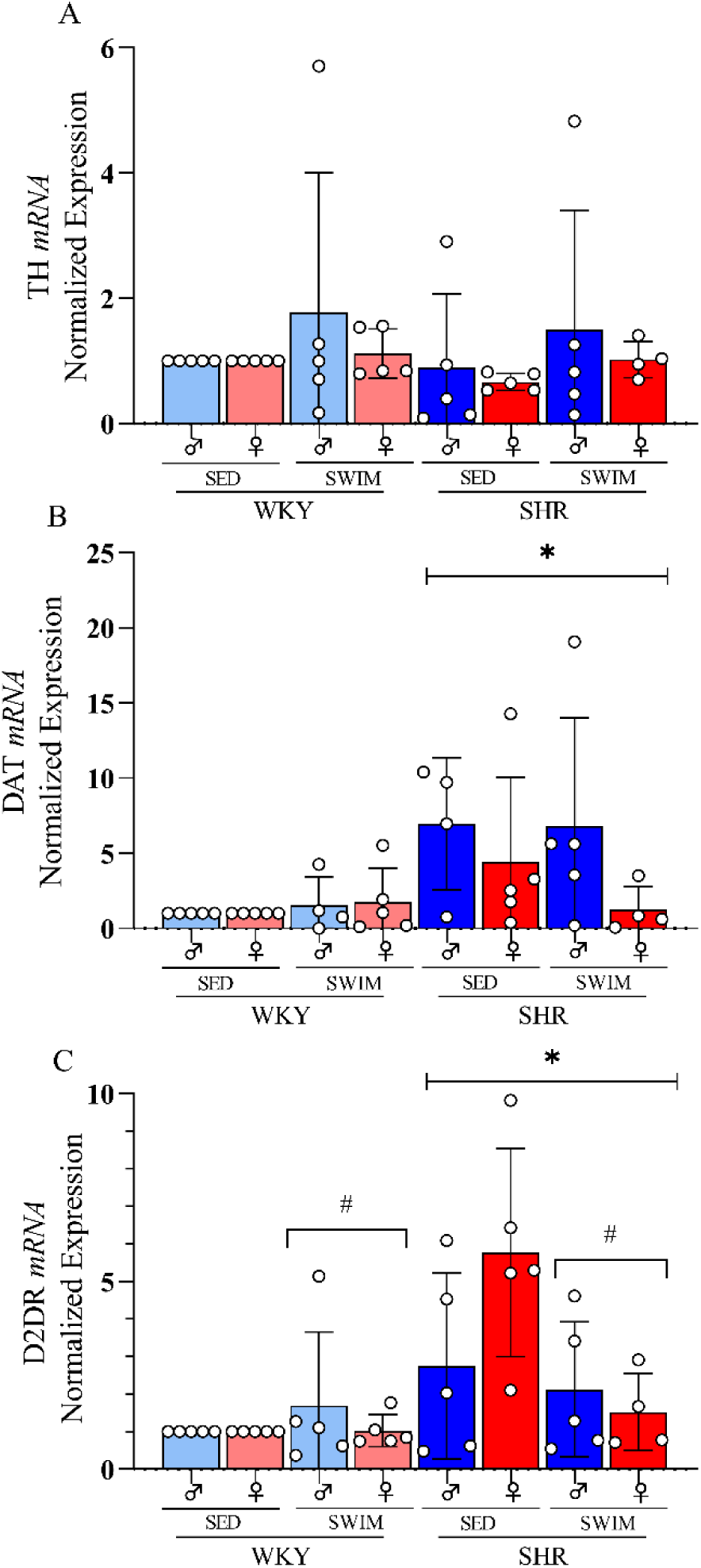
Effects of maternal swimming on expression of dopaminergic parameters in prefrontal cortex. Histogram represent the means ± SEM (n=3-5 animals per group) of expression of (A) tyrosine hydroxylase enzyme (TH), (B) dopamine transporters (DAT) and (C) dopaminergic D_2_ receptors (D_2_DR) in samples of the prefrontal cortex of WKY and SHR rats exposed or not to maternal swimming (SWIM and SED). *p<0.05 WKY vs. SHR strains, regardless of treatment and sex; ^#^p<0.05 SED vs. SWIM treatments in the same strain, regardless of sex (three-way ANOVA; *post hoc* Tukey).

## 4 Discussion

The present study investigated the effects of gestational swimming on behavioral and dopaminergic parameters in the offspring of WKY and SHR rats, an animal model of ADHD. SHR exhibited neurodevelopmental delay and novelty-seeking behavior in the adolescence phase, as well as increased expression of DAT and D2R in the PFC. In the SHRs, maternal gestational swimming improved motor reflexes of the offspring on postnatal days 7 and 14, and decreased novelty seeking and D2R expression in the adolescence phase.

Physical exercise significantly improves the quality of life, and it is generally accepted that regular physical exercise plays a critical role in improving brain structure and function (acho que precisa de umas refs aqui). Exercise increases neurogenesis and neurotransmitter release, promotes neuronal maturation, and improves learning and memory (Jeong et al., 2014; Kim et al., 2007; Ko et al., 2013). Moreover, regular physical activity can improve impaired social behavior in children with ADHD and reduces hyperactivity (Majorek et al., 2004). However, the possible behavioral and neurobiological consequences to the offspring of pregnant women diagnosed with ADHD who practice regular physical exercise are still unknown. Given that the etiology of ADHD involves genetic factors in 80% of cases, we hypothesize that gestational swimming could prevent the development of ADHD symptoms in the offspring.

Regular physical exercise is recommended for women during pregnancy. It reduces maternal weight gain, prevents gestational diabetes, improves blood circulation, and has significant positive effects on fetal development (Amorim et al., 2007; Bungum et al., 2000; Polley et al., 2002). In the current study, SHR females showed lower gestational weight gain compared to WKY. In addition, SHR mothers who performed gestational swimming had a lower final weight compared to sedentary mothers. There was no significant difference between swimming and sedentary mothers in WKY rats.

There are few studies investigating neurodevelopment in ADHD animal models. Ferguson et al. (2003) demonstrated that SHR were slower in the righting reflex and negative geotaxis compared to WKY and Sprague-Dawley rats and associated this difference to maturational delays in individuals with ADHD. An important parameter of animal development is weight gain during the neonatal period. SHR have lower weights than control WKY rats (Ferguson et al., 2003a). The results of the present study confirmed this difference between strains on both PND 7 and PND 14. Gestational swimming influenced weight gain on PND 7, decreasing weight in WKY animals and increasing weight in SHR. In the righting reflex test, gestational swimming proved beneficial on both PND 7 and PND14 in SHR, improving righting times compared to the SHR SED group.

The SHRs was slower than the WKY strain on both days of testing, supporting the neurodevelopmental delay. In the motor coordination test of negative geotaxis, there was a significant difference between the groups, mainly because all SHRs (groups SED and SWIM) were unable to turn 180° on PND7 and reached the maximum time in the test (60 seconds). On PND 14, we observed an improvement in this motor impairment in SHR compared with PND 7; however, SHR still showed a significant difference from the WKY controls.

In the sensory reflex test of olfactory recognition, the differences between strains are also easily detected. SHRs showed sensory impairment on PND 7: animals in the SHR SED group (both males and females) reached the maximum test time. On PND 14, the time spent to discriminate maternal odor was shorter than on PND 7, but SHR were still slower than WKY rats. The SHR SWIM group did not reach the maximum test time in PND 7, but still had sensory impairments. On PND 14, the benefits of gestational swimming were observed in SHR females, which had a lower odor discrimination index than females in the SHR SED group. Sanches et al. (2017) showed that swimming during pregnancy improves developmental reflexes in a model of hypoxia-ischemia. Although neonatal hypoxia-ischemia is considered a risk factor for ADHD, the present study is the first to show that gestational swimming improves neurodevelopmental aspects in an animal model of ADHD.

Adolescence is characterized by behavioral traits such as novelty seeking, risky behavior, and high social interaction, which promote the acquisition of skills necessary for maturation and independence (Shulman et al., 2016; Steinberg, 2010). It is also considered a period of high vulnerability to psychiatric disorders (Botdorf et al., 2017; Paus et al., 2008) and substance abuse (Chassin et al., 2002; Willoughby et al., 2013). Approximately 75% of children diagnosed with ADHD are thought to continue to have symptoms into adolescence (Katragadda and Schubiner, 2007). As in individuals with ADHD, the SHRs exhibited risk-taking and novelty-seeking behavior, in addition to impaired habituation memory. Interestingly, offspring whose mothers were exercised showed a lower interest in new objects than the SED group. This is an interesting observation because these behaviors are closely related to substance abuse in ADHD individuals, especially during adolescence (Pandolfo et al., 2007).

Studies demonstrate that physical exercise affects neural maturation during the critical development period (Kim et al., 2007; Klein et al., 2020; Matté et al., 2021; Sanches et al., 2017a) and therefore causes persistent behavioral and dopaminergic changes in an ADHD animal model (Ko et al., 2013).

SHR have been validated as an animal model for ADHD as, in addition to behavioral similarity, they exhibit imbalance in frontostriatal dopaminergic circuits, namely lower dopamine release and high expression of DAT and D2R. Dopaminergic dysfunction may contribute to behavioral changes observed in SHR (Russell, 2003; Sagvolden, 2000b; Sagvolden and Sergeant, 1998; Sandberg, 2005). We found that swimming during pregnancy decreased the expression of the D2R receptor in the frontal cortex of SHR offspring. Normalization of D2R expression could lead to regulation of dopaminergic phasic transmission, thus increasing the availability of dopamine at the synaptic cleft and improving some behavioral parameters of ADHD.

In conclusion, current results showed that maternal swimming during pregnancy prevented neonatal development impairments, improved behavioral aspects during adolescence, and normalized dopamine D2 receptors in the frontal cortex of SHR rats, and animal model of ADHD. These findings suggest that maternal gestational swimming holds potential to be used as a lifestyle preventative approach for ADHD.

## 5 Acknowledgements

AT was funded by a fellowship from Coordenação de Aperfeiçoamento de Pessoal de Nível Superior (CAPES) for her PhD. PP and STF was supported by Fundação Carlos Chagas Filho de Amparo à Pesquisa do Estado Rio de Janeiro (FAPERJ 26/201.430/2021, SEI-260003/004673/2021) and Conselho Nacional de Desenvolvimento Científico e Tecnológico (CNPq 308050/2022-3). MVL was supported by Serrapilheira Institute (R-2012-37967), FAPERJ (202.744/2019 and 210.316/2022) and CNPq (311487/2019-0 and 309278/2022-8)

## 6 Author contributions

PP and AT designed the study. AT, DM, and PS carried out the behavioral experiments. STF and MVL supervised biochemistry experiments. AT and ASF performed the western blotting and RT-PCR. AT wrote the first draft of the manuscript. STF, ML and PP wrote the final manuscript. All authors contributed to and have approved the final manuscript.

---

**Table 1.**
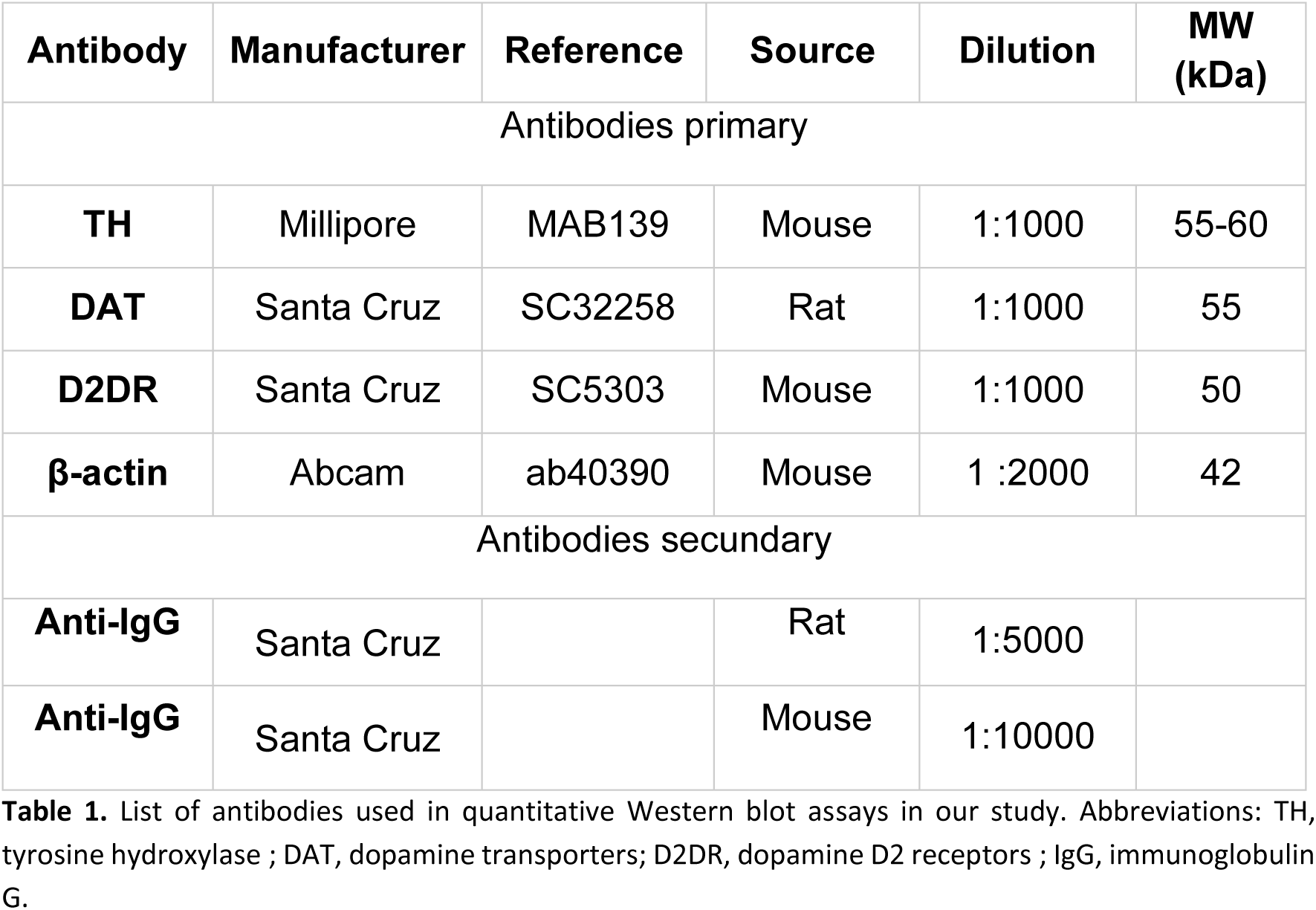
Antibodies used in the study.

List of antibodies used in quantitative Western blot assays in our study. Abbreviations: TH, tyrosine hydroxylase ; DAT, dopamine transporters; D2DR, dopamine D2 receptors ; IgG, immunoglobulin G.
| Antibody | Manufacturer | Reference | Source | Dilution | MW (kDa) |
| --- | --- | --- | --- | --- | --- |
| Antibodies primary |  |  |  |  |  |
| TH | Millipore | MAB139 | Mouse | 1:1000 | 55-60 |
| DAT | Santa Cruz | SC32258 | Rat | 1:1000 | 55 |
| D2DR | Santa Cruz | SC5303 | Mouse | 1:1000 | 50 |
| β-actin | Abcam | ab40390 | Mouse | 1 :2000 | 42 |
| Antibodies secondary |  |  |  |  |  |
| Anti-IgG | Santa Cruz |  | Rat | 1:5000 |  |
| Anti-IgG | Santa Cruz |  | Mouse | 1:10000 |  |

**Table 2.**
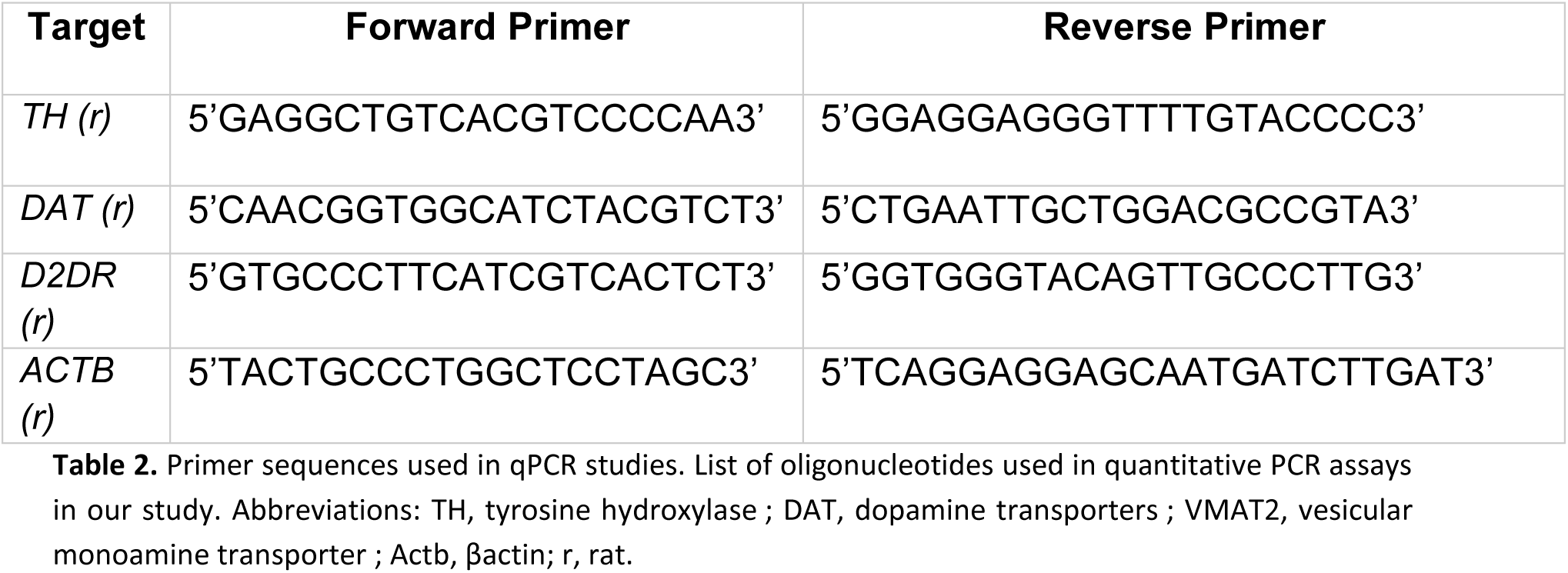
Primer sequences used in qPCR studies.

